# AnthOligo: Automating the design of oligonucleotides for capture/enrichment technologies

**DOI:** 10.1101/2019.12.12.873497

**Authors:** Pushkala Jayaraman, Timothy Mosbruger, Taishan Hu, Nikolaos G Tairis, Chao Wu, Peter M Clark, Monica D’Arcy, Deborah Ferriola, Katarzyna Mackiewicz, Xiaowu Gai, Dimitrios Monos, Mahdi Sarmady

**Affiliations:** Department of Pathology and Laboratory Medicine, The Children’s Hospital of Philadelphia, Philadelphia, PA USA; Division of Biomedical & Health Informatics, The Children’s Hospital of Philadelphia, Philadelphia, PA USA; Perelman School of Medicine, University of Pennsylvania. Philadelphia, PA USA

**Keywords:** region-specific extraction, oligo, primer design, enrichment, next-generation sequencing

## Abstract

**Summary:** A number of methods have been devised to address the need for targeted genomic resequencing. One of these methods, Region-specific extraction (RSE) of DNA is characterized by the capture of long DNA fragments (15-20 kb) by magnetic beads, after enzymatic extension of oligonucleotides hybridized to selected genomic regions. Facilitating the selection of the most optimal capture oligos targeting a region of interest, satisfying the properties of temperature (Tm) and entropy (ΔG), while minimizing the formation of primer dimers in a pooled experiment is therefore necessary. Manual design and selection of oligos becomes an extremely arduous task complicated by factors such as length of the target region and number of targeted regions. Here we describe, AnthOligo, a web-based application developed to optimally automate the process of generation of oligo sequences to be used for the targeting and capturing the continuum of large and complex genomic regions. Apart from generating oligos for RSE, this program may have wider applications in the design of customizable internal oligos to be used as baits for gene panel analysis or even probes for large-scale comparative genomic hybridization (CGH) array processes.

**Implementation and Availability:** The application written in Java8 and run on Tomcat9 is a lightweight Java Spring MVC framework that provides the user with a simple interface to upload an input file in BED format and customize parameters for each task. A Redis-like *MapReduce* framework is implemented to run sub-tasks in parallel to optimize time and system resources alongside a ‘task-queuing’ system that runs submitted jobs as a server-side background daemon. The task of probe design in AnthOligo commences when a user uploads an input file and concludes with the generation of a result-set containing an optimal set of region-specific oligos.

AnthOligo is currently available as a public web application with URL: http://antholigo.chop.edu.

## Introduction

Massively parallel sequencing, in particular, short-read technologies such as Exome Sequencing have become important milestones in genomic diagnosis. Newer technologies[1-3], such as long-read sequencing using linked-read strategy from 10x genomics[4] and single-molecule real-time (SMRT) sequencing approach from PacBio[5] focus on improving coverage over complex genomic regions to achieve finer resolution over sequence and structural rearrangements. Combining such a sequencing approach with a low-cost targeted enrichment methodology provides significant benefits of economy, data management and analysis and generates a resultant “capture” data that is further enriched for one’s regions due to longer reads spanning gaps and complex repeat regions.

Region-specific extraction (RSE) of DNA is a solution-based technique for enrichment of defined genomic regions of interest. The method’s cost-effective target-enrichment approach allows longer sequence templates up to 20 kb and a uniform depth of coverage across a region of interest.

Probe design for targeted enrichment is a requirement for any NGS test development. Although there exist many stand-alone tools and web-applications to help address requirements for varied target enrichment approaches, none can be implemented directly for the RSE method[8-17]. The advantage of this specific oligonucleotide design method is the ability to “space” the oligos evenly at a certain distance (thousands of bases) and thus achieve equivalent target specificity with fewer probes required as compared to the tiling approach (1X or 2X tiling density) by many custom “kit” provisions. Prior to automation of oligonucleotide design for capture/enrichment, an analyst would have to painstakingly filter the oligonucleotides to create sets of oligos manually by scanning a large matrix of dimer-dimer interactions. The task could exponentially increase in complexity and time when factors such as target region, size or number of regions increased. By streamlining the process of oligo design via an automated, statistically-motivated downstream processing algorithm [9, 14, 16], we estimate the tool saves man-hours by at least 10-fold. Here, we present AnthOligo, an automated application to design evenly-spaced capture oligos when provided with coordinates for genome-specific regions of interest. We have successfully implemented AnthOligo to design optimal capture oligos for the Zebrafish genomes[7] and additionally targeted and captured 4 MB section of the highly complex, MHC region in the human genome[6] in a solution-based capture. Most recently, additional sets of oligos have been designed, enriching the MHC by including publicly available MHC reference sequences from other cell lines that were either, partially known or fully completed [18]. The newest set of oligos have been successfully used in our new study (Manuscript in preparation).

### Implementation

**Step 1:** A region in the input file can range from a single exon to multiple megabases. A sliding window approach spanning 2kb overlapping every 100bp ensured thorough coverage of the region (Figure 1). Primer3[19] was used to generate internal oligos within each window using a repeat-masked reference sequence[20]. UCSC BLAT[21] was used to inspect sequence specificity across the oligos at a percentage identity threshold customized at 95%. The ‘susceptibility’ to form hairpins and duplexes was estimated by measuring their Tm and ΔG predictions by MFold[22] and UNAFold[23] for dimer stability based on the parameters of SantaLucia et al.[8, 10, 24, 25].

**Step 2:** For each region of interest, oligos that passed applicable thresholds from **Step1** were considered “candidates”. The algorithm modeled the storage of oligos and specific properties like ‘dimer interactions’ and ‘association by distance’ in a directed acyclic graph(DAG)[16] (Figure 1). For RSE method to be able to capture the entire region of interest (ROI), the first few “seed” oligos must lie within a short window across the start of the region. The graph object consisted of seed oligos or ‘root nodes’ and associated oligos became ‘child nodes’. Each ‘edge’ represented the user-defined distance between the root and child nodes. A depth-first-search (DFS) was then carried out to walk through “completed paths” in each graph. A path was “complete” when the “leaf” oligo was found within the end of the target region. Each completed path formed a “set” of oligos for the given region.

**Step 3:** Design of optimal collection of oligos for target capture using multiplex PCR required combinatorial optimization solutions[11, 17] (Figure 1). The number of heterodimer combinations C for *n* oligos for each input region could be calculated as:

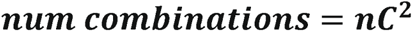

In order to get a resultant “*combination of set of oligos*” across all of the user-provided input regions, region-specific oligo sets were cross-compared across the input regions to ensure that oligos across regions did not dimerize with each other in solution. Every *m, k, p* number of oligos across M, K, P additional input regions increased this number of combinations somewhat exponentially:

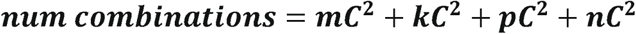

With increasing region size and number of regions, this became computationally intensive akin to the Np-complete ‘knapsack problem’. Heuristic optimization allowed for scalability without sacrificing quality of the capture design by returning the first available combination of oligo sets that satisfied our thresholds.

## Results and Discussion

Besides the published work (6,7), oligos have been designed for capturing several genomic regions associated with Noonan Syndrome (8 genes), Type 1 Diabetes (9 genomic regions), Crohn’s Disease (10 genomic regions) and retinitis pigmentosa (37 genomic regions) (available upon request). In each case the oligonucleotides performed well as observed by uniformity, sensitivity and average depth of coverage[6, 7]. To additionally validate the tool, the MHC of a random sample was captured and sequenced on the Illumina MiSeq. Alignment was performed using COX as reference, since the sample showed a closer match to COX than PGF reference. The average depth of coverage was estimated at 100x with 98.4% of positions >20X [Supplementary data Fig 1]. The reason we attempted another capture of the MHC region, besides the one published earlier (6), is because we needed to assess the success of the design using a random sample with unknown MHC sequence. The previously published capture (6) involved the PGF cell line, which has a known MHC sequence and the oligos were designed based on this known reference sequence. This time the Antholigo using a number of different reference MHC sequences (18) was used to generate a new set of oligos that presumably can target the MHC of any random DNA sample.

To capture sequence with acceptable range of accuracy and uniform representation across all the regions in multiplexed reactions, oligonucleotides must meet certain specifications in terms of sequence specificity, efficient oligo design with minimal interaction between the probes and optimal process time[14, 16, 17, 26]. AnthOligo was implemented to satisfy these requirements with the RSE method. It is well-understood that target capture design for multiplexed reactions is an NP-complete problem [14, 27]. Heuristic optimization was necessary to process large regions, upwards of 1Mb while identifying sets of evenly spaced capture oligonucleotides throughout the target region with target specificity[28]. Combinatorial approaches along with *MapReduce* framework helped multi-thread memory-intensive and data-intensive tasks to run within an optimal time.

Sequence specificity is governed by multiple factors, the majority of which are repeats in the genome and the presence of pseudogenes [10, 11, 29-32]. AnthOligo’s use of hard-masked reference file for generating oligos resolves this by avoiding possible repeat regions in the sequence. BLAT results were filtered by focusing on the specificity of the 3’ subsequence[33].

Although AnthOligo was developed to support the RSE method, its current abilities and flexibility for future enhancements may have wider applications in designing internal oligos that can be used to target the MHC using CRISPR-Cas9, baits for gene panel analysis or even probes for CgH array processes. AnthOligo is thus, a unique tool to an unaddressed domain and results show that it achieves the desired objectives.

## Acknowledgements

Thanks to Juan Carlos Perin for a great name, Dr. Kajia Cao, Dr. Chao Wu for help with the NP-Complete optimization problem.

## Funding

The project described was supported by Award Number P30DK019525 from the National Institute of Diabetes and Digestive and Kidney Diseases to DM.

---

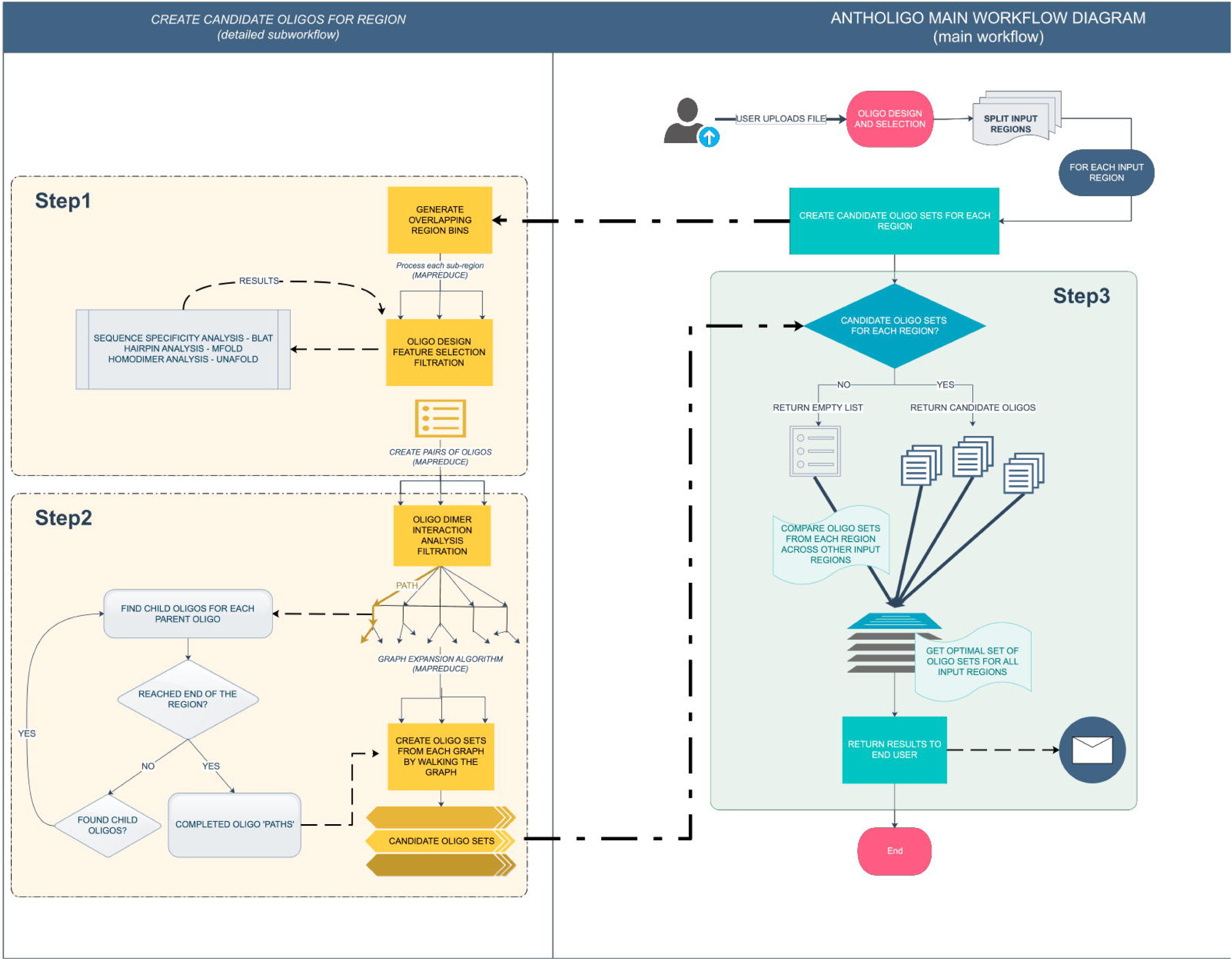

