## Supplementary data for "AnthOligo: Automating the design of oligonucleotides for capture/enrichment technologies"

**Algorithm Implementation (Supplementary)**

The **Help** page provides users with necessary information and a test file to begin using the application.

**Oligo design and Selection:** Primer3 showed great reproducibility over a window span and was preferential to certain regions. It can be theorized that the number of windows correlates linearly with region size. Given a region size, the number of oligos *n* that one can expect for a given target is:

$$\boldsymbol{RegionSize = 2000}\boldsymbol{n-100}\left( \boldsymbol{n-1} \right)\boldsymbol{+1}$$

Multiple smaller regions were processed faster than a single large region and hence, a really large region could be processed relatively faster when broken down into smaller regions. Performance was not compromised due to implementation of *MapReduce* and there was distinct advantage of increase in the oligo density distribution where regions of interest overlap.

**Hairpin and Oligo Dimer Structures:** Multiple oligo analysis tools[1-4] contain modules that predict the susceptibility of an oligo to form these structures.

###### **Creating optimal sets of oligo sets across regions:**

When designing targets for a multiplex PCR, it is customary to pool together multiple regions of interest in a single reaction. In order to group optimal oligo sets together, it is imperative to test dimer-dimer interactions of oligos across the sets so as to ensure optimal specificity during hybridization and to reasonably amplify all regions.

Increasing number of input regions and variability in region-sizes exponentially escalates number of oligo sets which in turn increases processing time and CPU power. The heuristic optimization generates the final result as the first available group of oligo sets instead of the best available result set[5, 6]. This allows for scalability without sacrificing quality of the capture design. On creation of a final set of oligos across all the input regions, an email is sent to the end user with results attached.

**Results and Discussion (Supplementary)**

Alignment was performed on the sequenced data using the following protocol:

$\#create index$

$\sim/applications/bwa-0.7.15/bwa index hg38\_cox.fa$

$\#map reads$

$\sim/applications/bwa-0.7.15/bwa mem -t 20 hg38\_cox.fa DM\_S1\_L001\_R1\_001.fastq.gz DM\_S1\_L001\_R2\_001.fastq.gz | samtools view -Sbh | samtools sort - -o DM\_10X.bam \&\& samtools index DM\_10X.bam$

$\#determine aligned (primary)$

$samtools view -c -F 2308 DM\_10X.bam$

$\#determine cox (primary)$

$samtools view -c -F 2308 DM\_10X\_cox.bam chr6\_GL000251v2\_alt:1143400-4734510$

$\#coverage$

$\sim/applications/bedtools2/bin/coverageBed -a region.bed -abam DM\_10X\_cox\_bwa.bam -d > DM\_cox\_bwa\_coverage.txt$

$\#bedgraph$

$\sim/applications/bedtools2/bin/genomeCoverageBed -ibam DM\_10X\_cox\_bwa.bam -bg > DM\_10X\_cox\_bwa.bg$

$\#bigwig$

$\sim/applications/bedGraphToBigWig DM\_10X\_cox\_bwa.bg ../../20190116\_10X/reference/hg38\_cox.fa.fai DM\_10X\_cox\_bwa.bw$

The human MHC reference is represented by many alternative cell lines, available in the UCSC database as standalone *hap_* sequences, while the hg19 human reference MHC region haplotype sequence is from PGF. For the oligos to clearly capture the diversity in the MHC class II region, these alternative cell line references may be a better match to some samples than PGF. Likewise, our sample was a closer match to COX reference per previously available haplotyping results. Therefore, coordinates for cell lines from 7 additional MHC reference sequences[18] were added to the input file to boost the enrichment, thus making the process somewhat “reference-inclusive”.

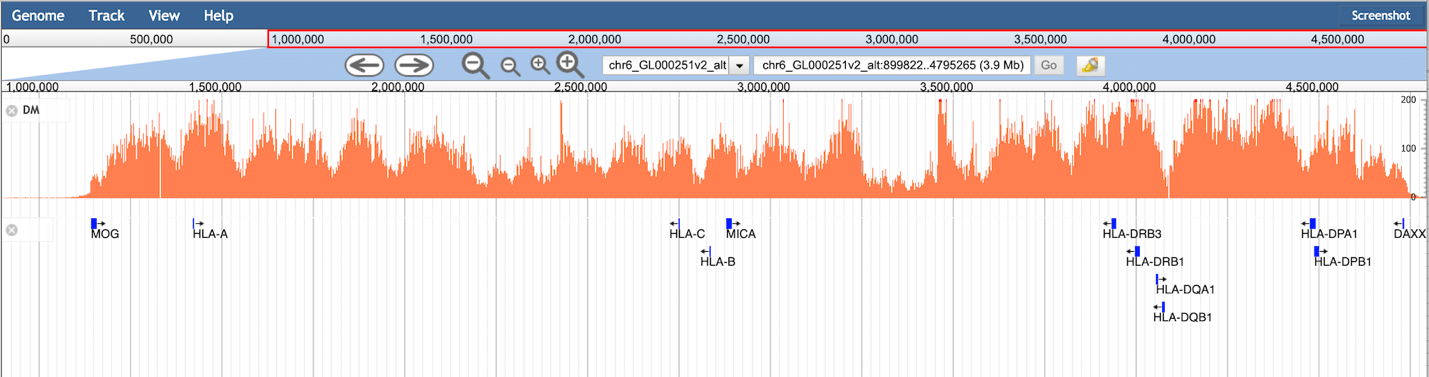

**Fig 1:** Capture coverage between MOG and DAXX for DM sample shows average coverage 100X with 98.4% bases >20X.

The application lists a limited number of genomes that it can generate oligos for. Additional genomes can easily be incorporated into the application upon request.

The publicly available version of AnthOligo does have a limitation on the cumulative size of all target regions provided by the end user and will notify the end user when that limit is exceeded in the file (cumulative region limited at 1MB). This is only because of the physical restrictions of resources on our production environment. However, if a user wishes to submit regions whose cumulative size is larger than this provided limit, we do have an internal version of AnthOligo with more robust computing power.
